# Checkpoint regulation of nuclear Tos4 defines S phase arrest in fission yeast

**DOI:** 10.1101/775130

**Authors:** Seong M. Kim, Vishnu P. Tripathi, Kuo-Fang Shen, Susan L. Forsburg

## Abstract

From yeast to humans, the cell cycle is tightly controlled by regulatory networks that regulate cell proliferation and can be monitored by dynamic visual markers in living cells. We have observed S phase progression by monitoring nuclear accumulation of the FHA-containing DNA binding protein Tos4, which is expressed in the G1/S phase transition. We use Tos4 localization to distinguish three classes of DNA replication mutants: those that arrest with an apparent 1C DNA content and accumulate Tos4 at the restrictive temperature; those that arrest with an apparent 2C DNA content, that do not accumulate Tos4; and those that proceed into mitosis despite a 1C DNA content, again without Tos4 accumulation. Our data indicate that Tos4 localization in these conditions is responsive to checkpoint kinases, with activation of the Cds1 checkpoint kinase promoting Tos4 retention in the nucleus, and activation of the Chk1 damage checkpoint promoting its turnover. Tos4 localization therefore allows us to monitor checkpoint-dependent activation that responds to replication failure in early versus late S phase.

## INTRODUCTION

The cell cycle proceeds through a rhythmic pattern of oscillators driven by cell-cycle specific transcription, patterns of protein modification, and protein degradation (reviewed in (Bertoli *et al.* 2013; Malumbres 2014; Alber *et al.* 2019)). Fission yeast is an important model system for studying cell cycle dynamics and genome stability. The rod-shaped cells are divided by medial fission with distinct cell morphologies (Piel and Tran 2009). Typically, mitosis is completed and S phase begins when cells are in a binucleate stage, prior to septation (Gomez and Forsburg 2004; Peng *et al.* 2005; Piel and Tran 2009). Thus, new-born cells are considered to be in late S to G2 phase, while S phase begins in binucleates (MacNeill and Fantes 1997). Distinguishing early from late S phase is typically done by monitoring nuclear DNA content by methods such as FACS or BrdU (Hodson *et al.* 2003; Sabatinos and Forsburg 2015b). Isotopic labeling methods suggest that the bulk of DNA synthesis is complete in a short time, leading to the conclusion that S phase is quite short and G2 phase extended (Nasmyth *et al.* 1979).

However, many replication mutants in fission yeast show an approximately 2C DNA content upon cell cycle arrest; based on genetic studies, this has been proposed to be late S phase (e.g., (Nurse *et al.* 1976; Nasmyth and Nurse 1981; Coxon *et al.* 1992; Forsburg and Nurse 1994)). Whether this arrest represents failure to duplicate specific late regions remains to be seen. Generally, late-replicating genome regions show increased prevalence of mutations and fragile sites (Beau *et al.* 1998; Stamatoyannopoulos *et al.* 2009; Lang and Murray 2011). Very late DNA replication has been observed, even into M phase for repair synthesis (Idrow *et al.* 1998; Bergoglio *et al.* 2013; Minocherhomji *et al.* 2015). Indeed, models of replication stress increasingly suggest the issue is not within early S phase but disruptions of chromosome segregation during mitosis (Zeman and Cimprich 2014; Minocherhomji *et al.* 2015; Ning *et al.* 2019).

We are interested in identifying early S phase cells and distinguishing them from late S phase or G2. Recent advances in live cell imaging have been accompanied by developing markers that are specific to particular cell cycle compartments. For example, the FUCCI (Fluorescent Ubiquitination-based Cell Cycle Indicator) system has been deployed using tagged, ubiquitylated proteins that are specific to G1/S or G2 cells (Sakaue-Sawano *et al.* 2008) and further refined by additional markers specific to G0 (Oki *et al.* 2014) or to multiple cell cycle phases (Bajar *et al.* 2016). These proteins vary temporally and spatially, giving a snapshot of cells in a particular cell cycle phase. There are excellent markers for mitotic landmarks including fluorescently tagged spindle pole body protein Sad1 (King and Drivas 2008) or tubulin (Sawin and Tran 2006), and septation is easily examined under light microscopy (Minet *et al.* 1979). We have developed and employed numerous tools to identify and characterize features of DNA synthesis and replication stress, including fluorescent tagged RPA and Rad52 proteins (Sabatinos *et al.* 2012, 2015; Green *et al.* 2015; Sabatinos and Forsburg 2015a; Escorcia and Forsburg 2017), and have also examined abnormal mitotic divisions in response to replication stress including nuclear envelope, cell membrane, and histone markers (Sabatinos *et al.* 2012, 2015; Escorcia and Forsburg 2017).

The forkhead-associated domain (FHA)-containing DNA binding protein Tos4 is conserved in budding and fission yeast (Kiang *et al.* 2009; Oliveira *et al.* 2012; Smolka *et al.* 2012). It is known to be regulated by the G1/S phase master transcription factor MBF (MluI-binding factor transcriptional complex) in both yeasts (Kiang *et al.* 2009; Oliveira *et al.* 2012). Tos4-GFP shows periodic accumulation in the nucleus coincident with S phase, consistent with its known regulation and the maturation timing of GFP (Kiang *et al.* 2009; Oliveira *et al.* 2012; Escorcia *et al.* 2019; Shen and Forsburg 2019). In fission yeast, it has been used in studies of cyclical re-replication induced by cyclin inhibition (Kiang *et al.* 2010). In this study, we characterize Tos4-GFP as a dynamic marker for S phase and determine its response to a variety of replication stresses, using both fluorescence microscopy and flow cytometry. We observe consistent timing of Tos4 accumulation relative to SPB duplication and septation in wild type cells. Tos4 persists in the nucleus of cells arrested in S phase by hydroxyurea (HU) or cell cycle mutant *cdc22-M45*, treatments which activate the replication checkpoint kinase Cds1. Consistent with this, accumulation of nuclear Tos4 requires Cds1 and specifically the kinase’s FHA domain. Surprisingly, however, replication mutants that show apparent late S phase arrest lack nuclear Tos4. This suggests that Tos4 specifically delineates an early stage of S phase and leads to the possibility that “late S phase” defined by replication mutants overlaps with what we commonly call G2 phase in which low yet detectable levels of DNA synthesis is occurring (Kelly and Callegari 2019).

## MATERIALS AND METHODS

### Yeast strains and media

*S. pombe* strains (Table 1) were grown in supplemented Edinburgh minimal medium (EMM) for live cell imaging, Western blot, and flow cytometry. Cells were treated with12 mM hydroxyurea (HU, Sigma), incubated at 36°C for 4h, or pre-treated with 12 mM HU for 2 h at 25°C and then incubated at 36°C for 4h.

**Table 1.** Yeast strains used in this study

| Strain | Genotype | Source |
| --- | --- | --- |
| FY8222 | *h- Tos4-GFP::KanMX6 Sad1+::DsRed-LEU2 leu1-32 ura4-D18 ade6-M210 can1-1* | Our stock |
| FY8678 | *h- cdc21-M68 Tos4-GFP::KanMX6 Sad1+::DsRed-LEU2 ura4-D18 leu1-32 ade6-(M216 or 704)* | Our stock |
| FY8682 | *h+ mcm4 (cdc21-m68)-ts-dg::ura4+ Tos4-GFP::KanMX6 Sad1+::DsRed-LEU2 ura4-D18 leu1-32* | Our stock |
| FY8851 | *h+ cdc25-22 Tos4-GFP::KanMX6 leu1-32 ura4-D18 ade6-M210* | Our stock |
| FY8853 | *h+ cdc10-V50 Tos4-GFP::KanMX6 Sad1+::DsRed-LEU2 ura4-D18 leu1-32* | Our stock |
| FY8855 | *h- cdc22-M45 Tos4-GFP::KanMX6 Sad1+::DsRed-LEU2 cdc22-M45 ura4-D18 ade6-M210 leu1-32* | Our stock |
| FY8939 | *h- nda3-KM311 Tos4-GFP::KanMX6 leu1-32 ura4-D18* | Our stock |
| FY9062 | *h- Δcds1::ura4+ Tos4-GFP::KanMX6 Sad1+::DsRed-LEU2 ura4-D18 leu1-32* | Our stock |
| FY9064 | *h- cdc21-c106::HphMx Tos4-GFP::KanMX6 Sad1+::DsRed-LEU2 ura4-D18 ade6-M210 leu1-32* | Our stock |
| FY9074 | *h+ cdc17-M75(kg) Tos4-GFP::KanMX6 Sad1+::DsRed-LEU2 ura4-D18 leu1-32 can1-1* | Our stock |
| FY9075 | *h- cdc17-K42(kg) Tos4-GFP::KanMX6 Sad1+::DsRed-LEU2 ura4-D18 leu1-32 ade6-M210 can1-1* | Our stock |
| FY9120 | *h- cut9-665 Tos4-GFP::KanMX6 sad1+::DsRed-LEU2 ura4-D18 leu1-32 ade6-M210* | Our stock |
| FY9126 | *h- Δcds1::ura4+ cdc21-M68 Tos4-GFP::KanMX6 Sad1+::DsRed-LEU2 ura4-D18 leu1-32* | Our stock |
| FY9127 | *h+ Δchk1::ura4+ cdc21-M68(mcm4)-dg::ura4+ Tos4-GFP::KanMX6 Sad1+::DsRed-LEU2 ura4-D18 leu1-32* | Our stock |
| FY9128 | *h+ cdc18 -K46 Tos4-GFP::KanMX6 Sad1+::DsRed-LEU2 leu1-32 ura4-D18 ade6-M210* | Our stock |
| FY9129 | *h- Δcds1::ura4+ cdc22-M45 Tos4-GFP::KanMX6 Sad1+::DsRed-LEU2 ura4-D18 leu1-32* | Our stock |
| FY9131 | *h- cdc6-23 Tos4-GFP::KanMX6 Sad1+::DsRed-LEU2 ura4-D18 leu1-32* | Our stock |
| FY9133 | *h- pold-ts2 (cdc6-ts2) Tos4-GFP::KanMX6 Sad1+::DsRed-LEU2 ura4-D18 leu1-32* | Our stock |
| FY9135 | *h+ cdc27-K3 Tos4-GFP::KanMX6 Sad1+::DsRed-LEU2 ura4-D18 leu1-32 can1-1* | Our stock |
| FY9157 | *h- rad4-116 Tos4-GFP::KanMX6 Sad1+::DsRed-LEU2 ura4-D18 leu1-32 ade6-M210 can1-1* | Our stock |
| FY9158 | *h+ hsk1-1312 Tos4-GFP::KanMX6 Sad1+::DsRed-LEU2 ura4-D18 leu1-32 ade6-M210 his3-D1* | Our stock |
| FY9159 | *h+ cds1-FHA\*:2HA6His:ura4+:leu1+ Tos4-GFP::KanMX6 leu1-32 ura4-D18* | P. Russell |
| FY9180 | *h- goa1-U53 (sna41ts) Tos4-GFP::KanMX6 Sad1+::DsRed-LEU2 leu1-32 ura4-D18 his3-D1 ade6-M210* | Our stock |
| FY9283 | *h+ cut4-533 Tos4-GFP::KanMX6 Sad1+::DsRed-LEU2 leu1-32* | Our stock |
| FY9284 | *h+ nuc2-663 Tos4-GFP::KanMX6 Sad1+::DsRed-LEU2 leu1-32 ura4-D18* | Our stock |

### Live-cell microscopy

Cells cultured in supplemented EMM media were placed on 2% agarose pads sealed with VaLaP (1/1/1 [wt/wt/wt] Vaseline/lanolin/paraffin) for live-cell imaging. Images were acquired using a DeltaVision microscope (with softWoRx version 4.1; GE, Issaquah, WA) using a 60x (NA 1.4 PlanApo) lens, solid-state illuminator, and 12-bit CCD camera. Images were deconvolved and maximum intensity projected for fluorescence images (sofrWoRX) and transmitted light images were inverted and added for outline of the cells (ImageJ) (Schindelin *et al.* 2012).

### Western blot

Proteins extracts were prepared from equal number of Tos4-GFP cells in asynchronous culture grown in supplemented EMM media, after treatment with 12 mM hydroxyurea (HU), and after washing twice with media for release from HU. Cells in mid-log phase were harvested and whole-cell protein extract was prepared by vortexing acid-washed glass beads in 20% trichloroacetic acid (TCA) and washing beads with 5% TCA. Lysates were boiled for 5 min in Laemmli Sample buffer (4%SDS, 60 mM Tris-HCl, pH 6.8, 5% glycerol, 4% 2-mercaptoethanol, 0.01% bromophenol blue) and analyzed by 4-12% SDS-PAGE (Expedeon), followed by immunoblotting with rabbit anti-GFP (Abcam 290; 1:1000) and rabbit anti-cdc2 (gift from Nurse lab; 1:4000) as loading control. After secondary antibody (anti-rabbit Alexa Flour 488; 1:4000) incubation, blots were developed using Amersham Typhoon biomolecular imager.

### Flow cytometry

Cells were fixed in cold 70% ethanol and processed in 50 mM sodium citrate, 100 μg/ml RNase A, and 8 µg/ml propidium iodide (PI). Samples were sonicated and then run on the flow cytometer (BD Accuri C6 Plus) for GFP and PI.

### Statistical Analysis

A two-tailed Student’s t-test was used to determine significance: * P < 0.05, ** P < 0.01, *** P < 0.001, n.s. not significant. Error bars represent Standard Error (SE).

### Data Availability Statement

Strains and plasmids are available upon request. The authors affirm that all data necessary for confirming the conclusions of the article are present within the article, figures, and tables.

## RESULTS

### Dynamics of Tos4 in cycling wild type cells

To observe the dynamics of Tos4 accumulation in relation to the cell cycle, we imaged cells with Tos4-GFP and the spindle pole marker Sad1-DsRed. Tos4 accumulated in the nucleus of dividing cells, correlating to cells early S phase (Figure 1A and Movie 1). Nuclear Tos4-GFP was present largely in binucleate cells, corresponding to early S phase, and was also observed in some short mononucleate cells (newborns) following completion of septation, but was absent as the cells elongated, suggesting that nuclear Tos4 is lost in late S or G2. The duration of the presence of nuclear Tos4-GFP was about 60 minutes (Figure 1B) which was about 18% of the time of cell cycle (Figure 1C). Timing of nuclear Tos4-GFP relative to spindle duplication varied little between individual cells. We next examined the use of Tos4 as a dynamic marker to characterize S phase.

**Figure 1.**
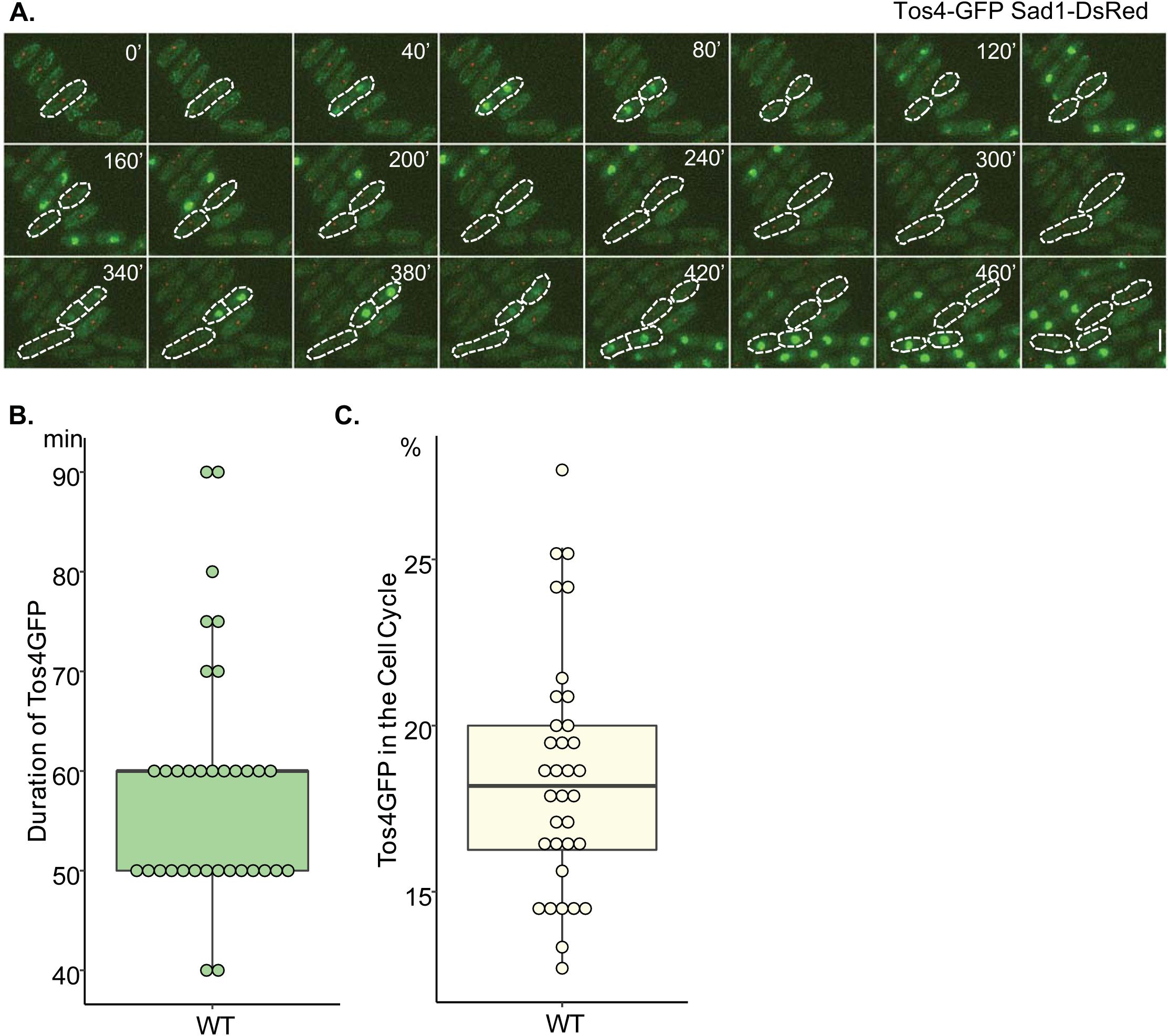
Tos4-GFP is present in nuclei of dividing cells. (**A**) Live cell imaging of WT cells containing Tos4-GFP and Sad1-DsRed was followed at 25°C for 8 hours. Nuclear Tos4-GFP signal is observed after duplication of SPBs and it dissappears from the nucleus before cells enter the next round of mitosis. Scale bar, 5 µm. (**B**) Duration of the presence of nuclear Tos4-GFP. The presence of nuclear Tos4-GFP was determined by measuring the nuclear Tos4-GFP signals using the ImageJ software (Schindelin *et al.* 2012). Nuclear Tos4-GFP signals was counted as a positive when nuclear Tos4-GFP signal is greater than 50 (scale 0-255) after the background subtraction using ImageJ. (**C**) Ratio of nuclear Tos4-GFP duration versus the cell cycle duration is presented. Duration of the cell cycle was determined by measurement of the timing between the first and second separation of SPBs. Sad1-DsRed was used to follow the separation of SPBs.

### Tos4 accumulation in cell cycle arrest defines early S phase

Treatment with hydroxyurea (HU) or temperature sensitive mutation of the ribonucleotide reductase component *cdc22* leads to depletion of nucleotide pools, and arrest of cells in early S phase, with a largely unreplicated DNA content (Timson 1975; Sarabia *et al.* 1993). We observed Tos4-GFP accumulation both in HU-treated or temperature-sensitive *cdc22-M45* at the restrictive temperature (Figure 2A,B). Tos4 was depleted when *cdc22-M45* cells were released back to the permissive temperature (25°C), consistent with return to the cell cycle (Figure 2B). Cells arrested in G1 by mutation of the MBF transcription factor that regulates *tos4*+ expression (*cdc10-V50*) or in G2 by the mitotic inducing phosphatase (*cdc25-22*) showed no nuclear accumulation of Tos4 at 36°C but gained nuclear Tos4 upon release to 25°C (Figure 2B). Similarly, cells arrested at mitosis (*nda3-KM311*) had no nuclear Tos4 at 17°C but gained nuclear Tos4 when released to 32°C (Figure 2C).

**Figure 2.**
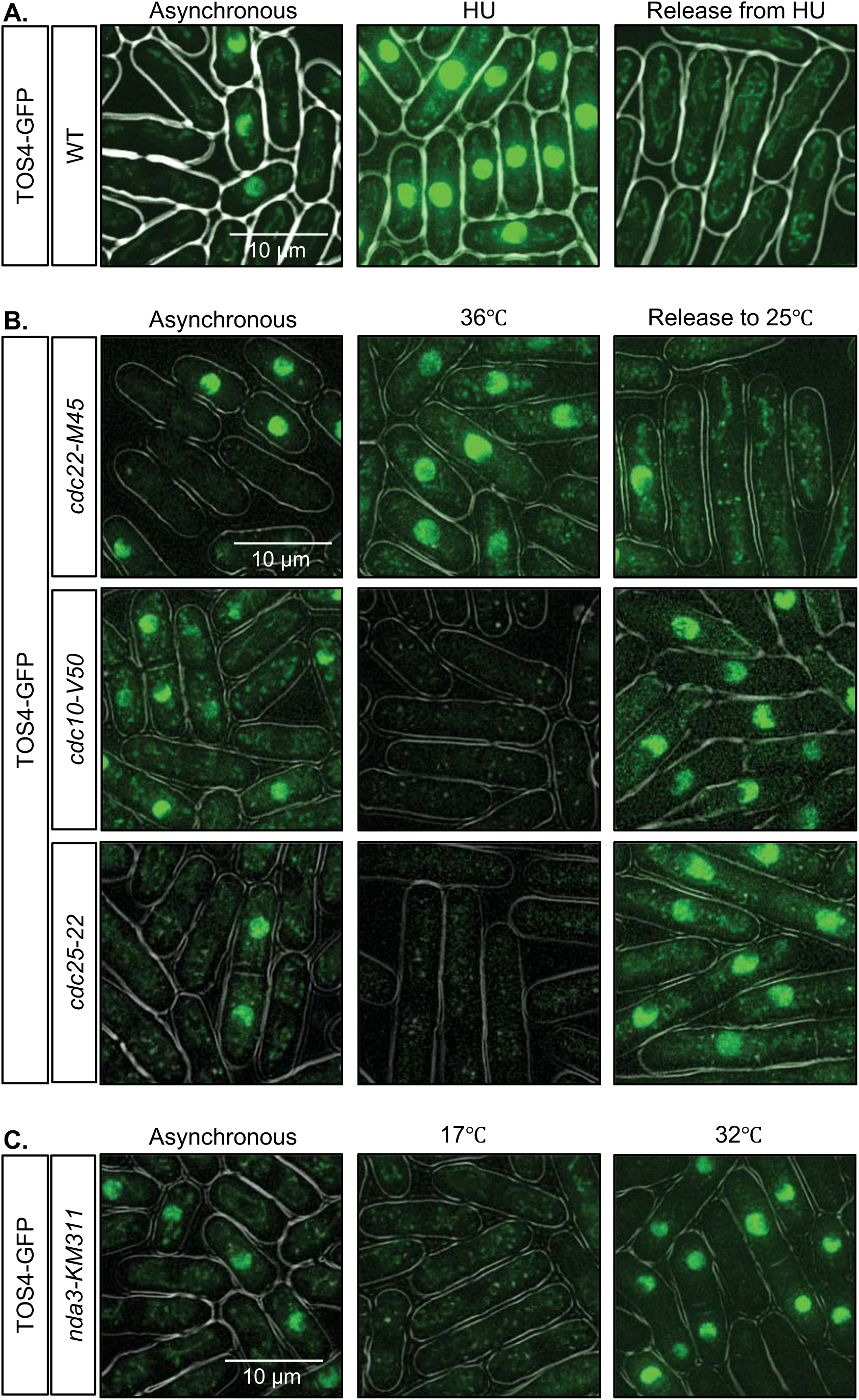
Tos4-GFP accumulate in the nuclei of cells arrested in early S phase. (**A**) WT cells were imaged for Tos4-GFP in asynchronous culture, after treatment with 12 mM HU for 4 h, and 1 h after release from HU. (**B**) Temperature-sensitive cell cycle mutants, *cdc22-M45* (S phase arrest), *cdc10-V50* (G1 phase arrest), *cdc25-22* (G2 phase arrest) were imaged for Tos4-GFP in asynchronous culture at 25°C, after 4 h at 36°C, or after 1 h-1.5 h after release to 25°C. (**C**) Cold-sensitive *nda3-KM311* (M phase arrest) was imaged for Tos4-GFP in in asynchronous culture at 32°C, after 4 h at 17°C, or after 0.5 h after release to 32°C.

Next, we examined Tos4 accumulation in a variety of S phase mutants. During replication, the MCM helicase, which comprises six subunits, unwinds the DNA duplex and promote replication initiation and progression (Forsburg 2004). The canonical temperature-sensitive mutant *mcm4-M68 (mcm4-ts)* synthesizes a near 2C DNA content at restrictive temperature (36°C) but shows low viability when released to permissive temperature (25°C) (Nasmyth and Nurse 1981; Coxon *et al.* 1992; Liang *et al.* 1999; Sabatinos *et al.* 2015). A large C-terminal truncation mutant *mcm4-c106*, also shows 2C DNA content at 36°C but much higher viability upon release than *mcm4-ts* (Nitani *et al.* 2008; Ranatunga and Forsburg 2016). A different temperature allele *mcm4-dg* that has a degron cassette added to *mcm4-ts*, undergoes rapid protein turnover at 36°C with limited DNA synthesis and a 1C DNA content (Lindner *et al.* 2002; Sabatinos *et al.* 2015) although it fails to arrest divisions (Sabatinos *et al.* 2015). Interestingly, all three *mcm4* mutants lacked nuclear Tos4 when placed at 36°C, even though they have different DNA contents and phenotypes (Figure 3A). We also tested temperature-sensitive mutants affecting the MCM loader *cdc18-K46*, DNA ligase mutants (*cdc17-M45* and *cdc17-K42*), and mutants affecting DNA polymerase delta subunits (*cdc6-23*, *cdc6-ts2*, and *cdc27-K3*). All of these arrest with a near 2C DNA content (Nasmyth and Nurse 1981). None maintained nuclear Tos4 (Figure 3B,C). Thus, Tos4 accumulation is different in early S phase (HU, *cdc22*) compared to late S phase mutants, and its accumulation is not limited by DNA content.

**Figure 3.**
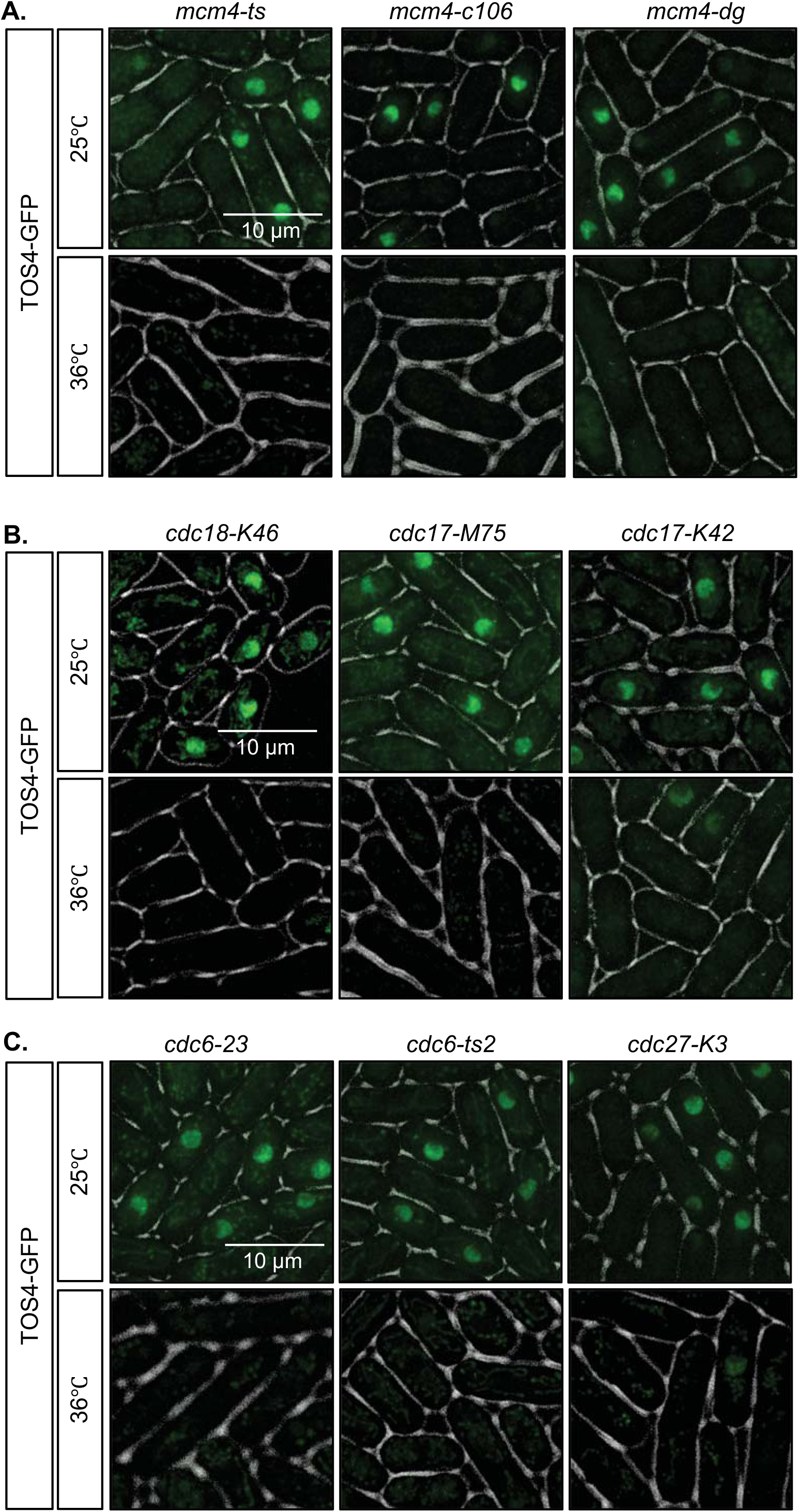
Replication mutants lack nuclear Tos4-GFP. Temperature-sensitive Mcm4 helicase mutants (*mcm4-ts, mcm4-c106, mcm4-dg*) (**A**), MCM loader mutant (*cdc18-K46*), ligase mutants (*cdc17-M45 and cdc17-K42*) (**B**), and polymerase delta mutants (*cdc6-23, cdc6-ts2, and cdc27-K3*) (**C**) were imaged for Tos4-GFP in asynchronous culture at 25°C or after 4 h at 36°C.

The Anaphase-Promoting Complex (APC) is an ubiquitin ligase that targets various proteins for proteasome-mediated degradation (Harper *et al.* 2002; Sivakumar and Gorbsky 2015). In budding yeast, Tos4 interacts with Cdh1, a WD40-repeat-containing activator of APC complex that recognizes degradation motifs in substrates (Ostapenko *et al.* 2012). Cdh1^Sc^ deletion results in partial stabilization of Tos4^Sc^ (Ostapenko *et al.* 2012) but not the temperature sensitive APC mutant *cdc23-1* ^Sc^, suggesting Tos4 protein turnover does not depend on APC in budding yeast. We observed Tos4-GFP in three different temperature sensitive APC mutants: *cut9-665*, *cut4-533*, and *nuc2-663*. At 36°C, Tos4 did not accumulate in any of these APC mutants (Figure 4A), consistent with a cell cycle arrest in mitosis. We pretreated APC mutants with HU and then released to 36°C. If Tos4 protein is a target for APC-mediated degradation, we reasoned Tos4 would remain nuclear. HU-treated APC mutant cells accumulated nuclear Tos4 but lost the signal when released to 36°C (Figure 4B). This suggests that similar to budding yeast, Tos4 is unlikely to be an APC target in fission yeast.

**Figure 4.**
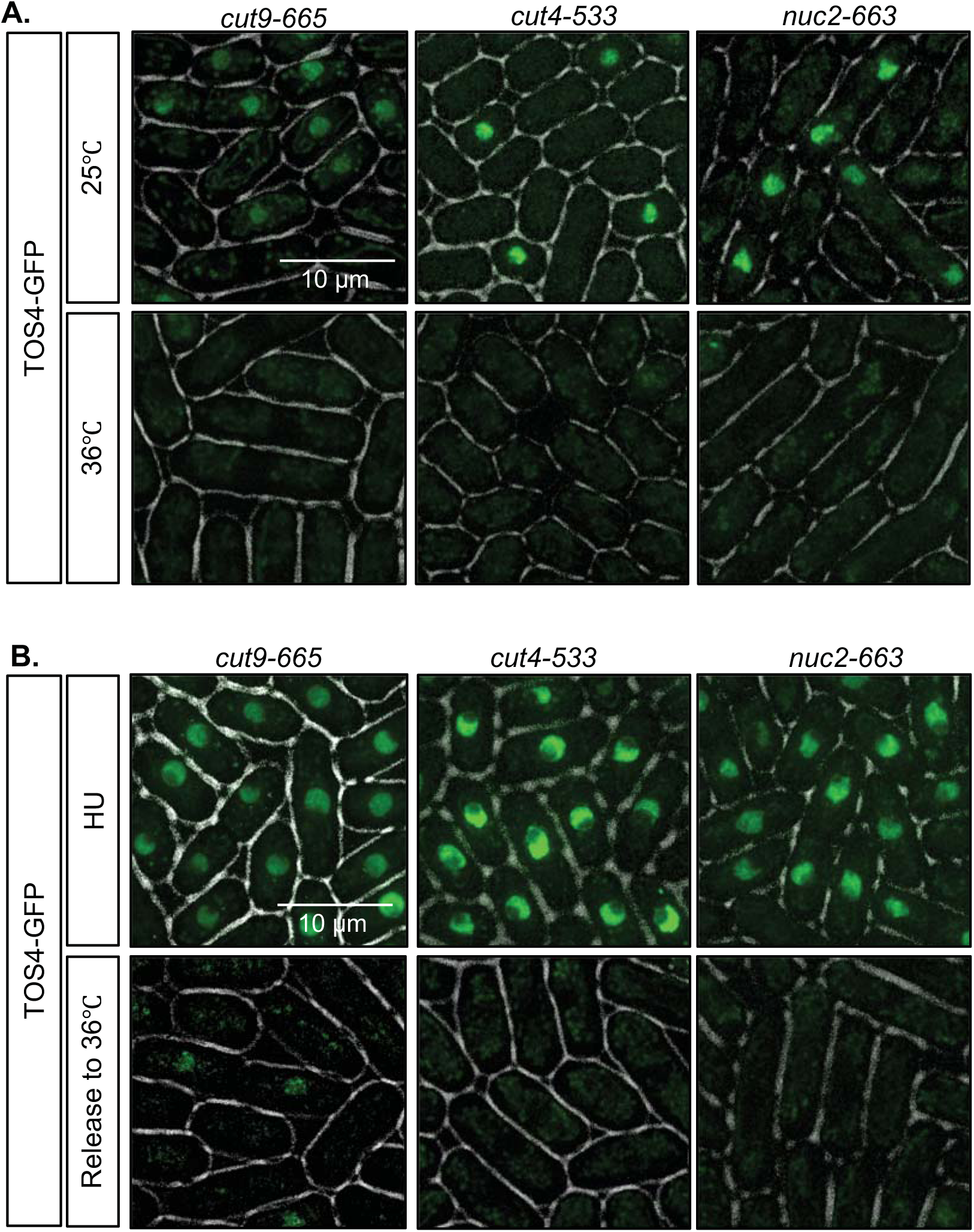
Tos4-GFP is not targeted for APC-mediated degradation. (**A**) Temperature-sensitive APC mutants (*cut9-665*, *cut4-533*, and *nuc2-663*) were imaged for Tos4-GFP in asynchronous culture at 25°C or after 4 h at 36°C. (**B**) APC mutants were imaged for Tos4-GFP after treatment with 12 mM HU for 4 h and 1 h after release from HU.

### Tos4 localization correlated with protein levels

We determined whether observed accumulation of nuclear Tos4 is due to nuclear localization of Tos4 alone or whether it correlates with protein levels changes during S phase, using western blot analysis and flow cytometry (FACS) analysis. Lysates were collected from cells arrested in S phase with HU and released. Tos4 protein level increased in cells arrested in S phase compared to cells in asynchronous culture (Figure 5A). Tos4 protein level decreased to basal levels 30-60 min after release from HU. We also used FACS analysis to detect the GFP signal. This showed similar results, with the GFP peak increased in cells arrested in S phase and decreased back as cells were released from HU (Figure 5B). These results demonstrate that both Tos4 localization and protein turnover are regulated during S phase.

**Figure 5.**
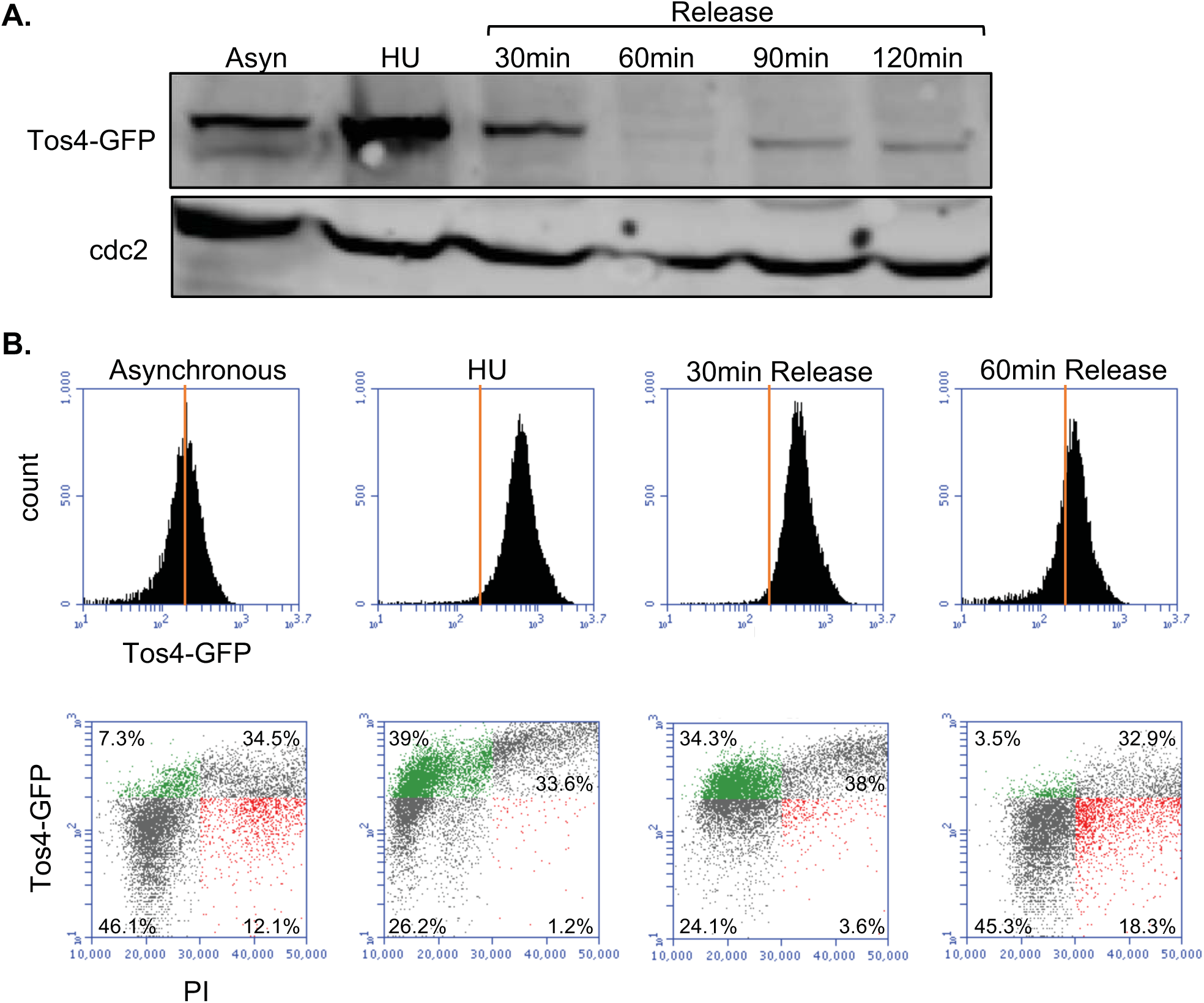
Tos4 protein level is increased in S phase cells arrested by HU. (**A**) WT cells with Tos4-GFP in asynchronous culture, after treatment with 12 mM HU, or after release from HU were lysed and immunoblotted for GFP and cdc2 (loading control). (**B**) WT cells used in (**A**) were fixed in 70% ethanol, and FACS analyzed for GFP and propidium iodide (PI). Green in scatter plot represents population with high GFP and low PI while red represents population with low GFP and high PI.

### Tos4 accumulation in early S phase is Cds1-dependent

HU blocks DNA synthesis by depleting deoxynucleoside triphosphate (dNTP) pools (Reichard 1988), which results in activation of the replication checkpoint kinase Cds1 (Lopes *et al.* 2001; Kai *et al.* 2005). Cds1 stabilizes replication forks and prevents cell division during replication arrest (Lindsay *et al.* 1998; Kai and Wang 2003). Previously, we showed that *cds1*∆ mutants fail to stop DNA synthesis during HU treatment, with lethal consequences (Sabatinos *et al.* 2012). Activation of Cds1 has been shown to upregulate the MBF transcription factor (de Bruin *et al.* 2008; Dutta *et al.* 2008; Chu *et al.* 2009; Oliveira *et al.* 2012) (reviewed in (Smolka *et al.* 2012; Bertoli *et al.* 2013)). Consistent with this, we observed that nuclear Tos4 accumulation during HU treatment or in *cdc22-M45* arrest is Cds1-dependent (Figure 6A,B). We also observed that this requires the forkhead-associated domain (FHA) of Cds1, a phospho-peptide-binding module that mediates association with proteins such as Mrc1 and Mus81 (Boddy *et al.* 2000; Tanaka and Russell 2004). The *cds1-fha\** allele has mutations at two highly conserved residues (S79A and H82A) in the FHA domain and decreases DNA damage tolerance (Boddy *et al.* 2000). Similar to *cds1*Δ, the *cds1-fha\** cells treated with HU did not show nuclear Tos4 (Figure 6A).

**Figure 6.**
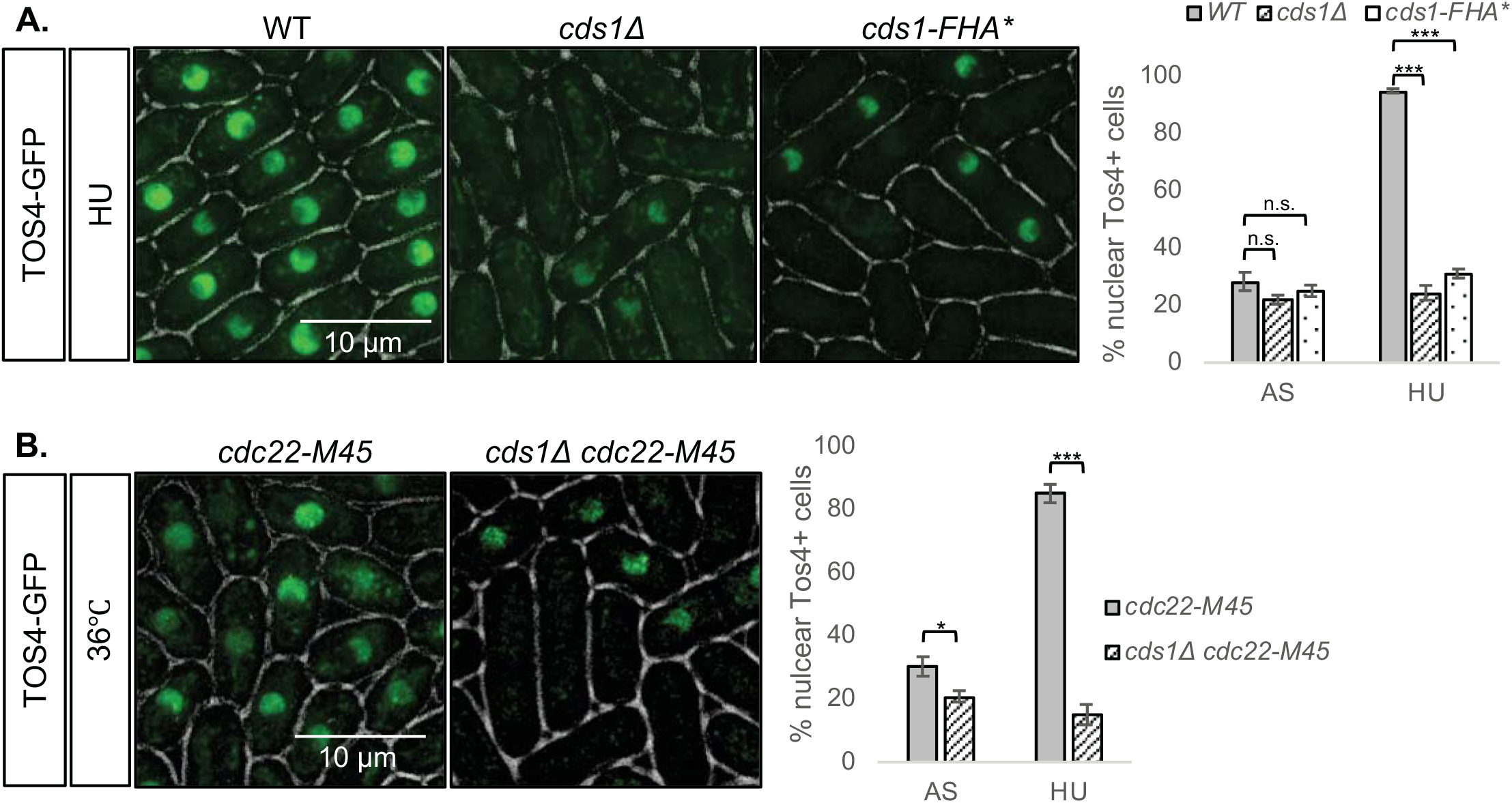
Cds1 is required for nuclear Tos4 accumulation in cells arrested in early S phase by HU or *cdc22-M45*. (**A**) *WT, cds1Δ, cds1-FHA\** cells were imaged for Tos4-GFP after treatment with 12 mM HU. Right, quantification of % cells with nuclear Tos4-GFP. (**B**) *cdc22-M45* and *cds1Δ cdc22-M45* cells were imaged for Tos4-GFP after 4 h at 36°C. Right, quantification of % cells with nuclear Tos4-GFP. A two-tailed Student’s t-test was used to determine significance: * P < 0.05, *** P < 0.001, n.s. not significant. Error bars represent Standard Error (SE).

We next examined whether activating Cds1 with HU first would be sufficient to maintain nuclear Tos4 in cells with temperature-sensitive mutations in replication mutants that normally do not accumulate Tos4. Temperature-sensitive *mcm4* mutants, *mcm4-ts* and *mcm4-dg*, treated with HU accumulated nuclear Tos4 as expected, but this was lost upon release from HU to 36°C (Figure 7A,B), demonstrating that transient hyperactivation of Cds1 by HU is not sufficient to maintain nuclear Tos4. We next asked whether there was a difference if we maintained HU treatment at the restrictive temperature, so we shifted *mcm4-ts* and *mcm4-dg* from HU at 25°C to HU at 36°C. The *mcm4-ts* cells maintain nuclear Tos4 under both temperature conditions, and this depends upon Cds1 (Figure 7A,B). Other replication mutants *cdc45/sna41, cdc18, cdc6, cdc17*, and *cdc27* also maintain nuclear Tos4 in the continued presence of HU at 36°C (Figure S1). Surprisingly, however, *mcm4-dg* cells do not maintain nuclear Tos4 in HU at 36°C (Figure 7A,B). Additionally, we showed that this loss of Tos4 in HU at 36°C in *mcm4-dg* is rescued by deletion of the damage checkpoint kinase Chk1 (Figure 7A,B).

**Figure 7.**
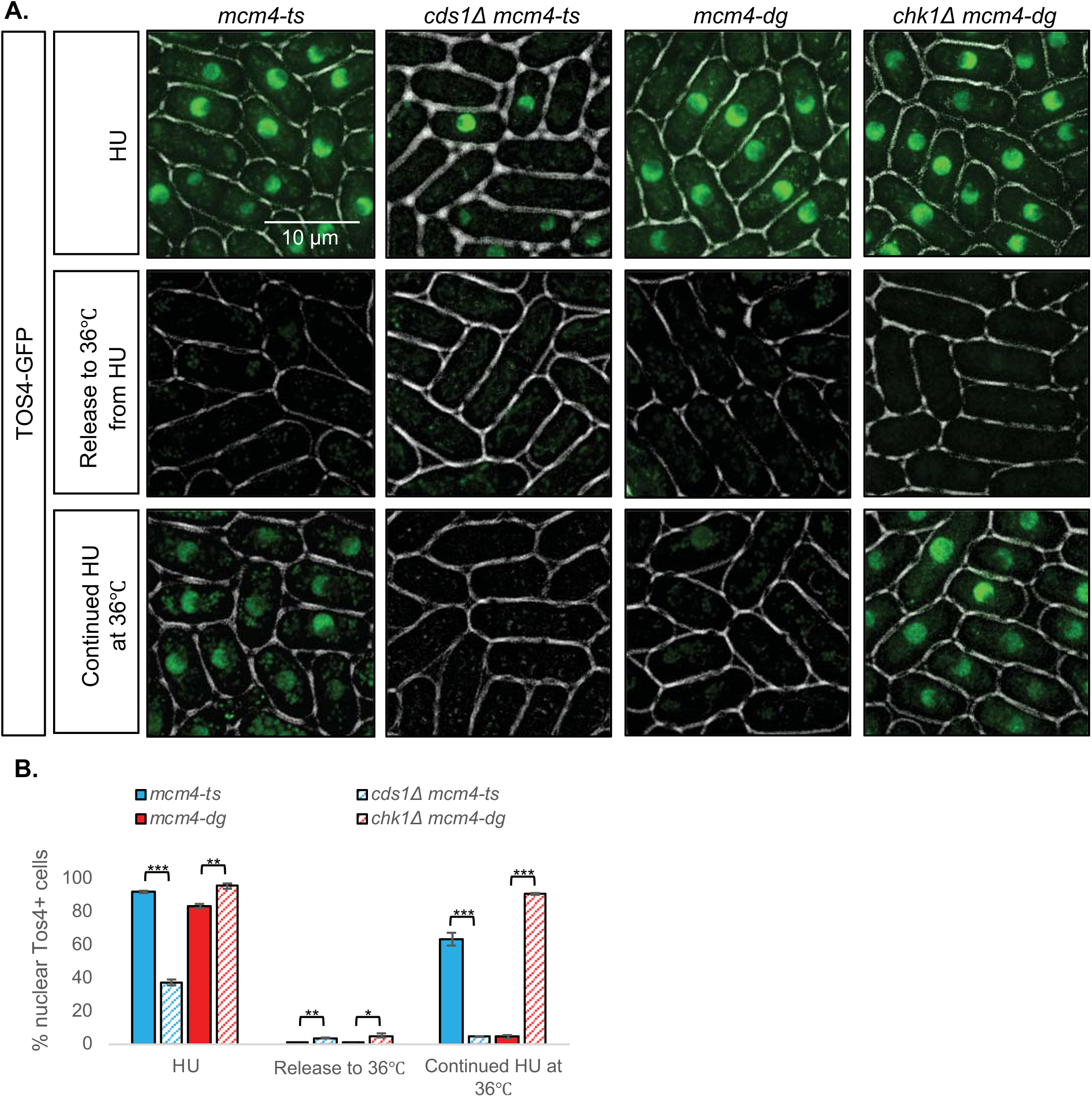
Continued Cds1 activation induces S phase arrest in *mcm4-ts* but not *mcm4-dg*. (**A**) *mcm4-ts, cds1Δ mcm4-ts, mcm4-dg, chk1Δ mcm4-dg* were imaged for Tos4-GFP after treatment with 12 mM HU at 25°C, after release from HU to 36°C, or after pre-treatment with HU at 25°C then transfer to 36°C. (**B**) Quantification of % cells with nuclear Tos4-GFP in (**A**). A two-tailed Student’s t-test was used to determine significance: * P < 0.05, ** P < 0.01, *** P < 0.001, Error bars represent Standard Error (SE).

The *mcm4-dg* allele is unusual as it bypasses normal cell cycle arrest and continues into mitosis despite the absence of substantial DNA synthesis (Sabatinos *et al.* 2015). We looked at two additional temperature sensitive replication mutants. The *hsk1*-1312 mutation affects, the catalytic subunit of the fission yeast Dbf4-dependent kinase (DDK) that regulates initiation of DNA replication via MCM, and *rad4-116 (cut5)* is also required for initiation, yet both proceed into mitosis at the restrictive temperature (Saka *et al.* 1994; McFarlane *et al.* 1997; Ostapenko *et al.* 2012). Similar to *mcm4-dg*, both these mutants lose nuclear Tos4 at 36°C and fail to maintain nuclear Tos4 in continued presence of HU at 36°C (Figure 8A).

**Figure 8.**
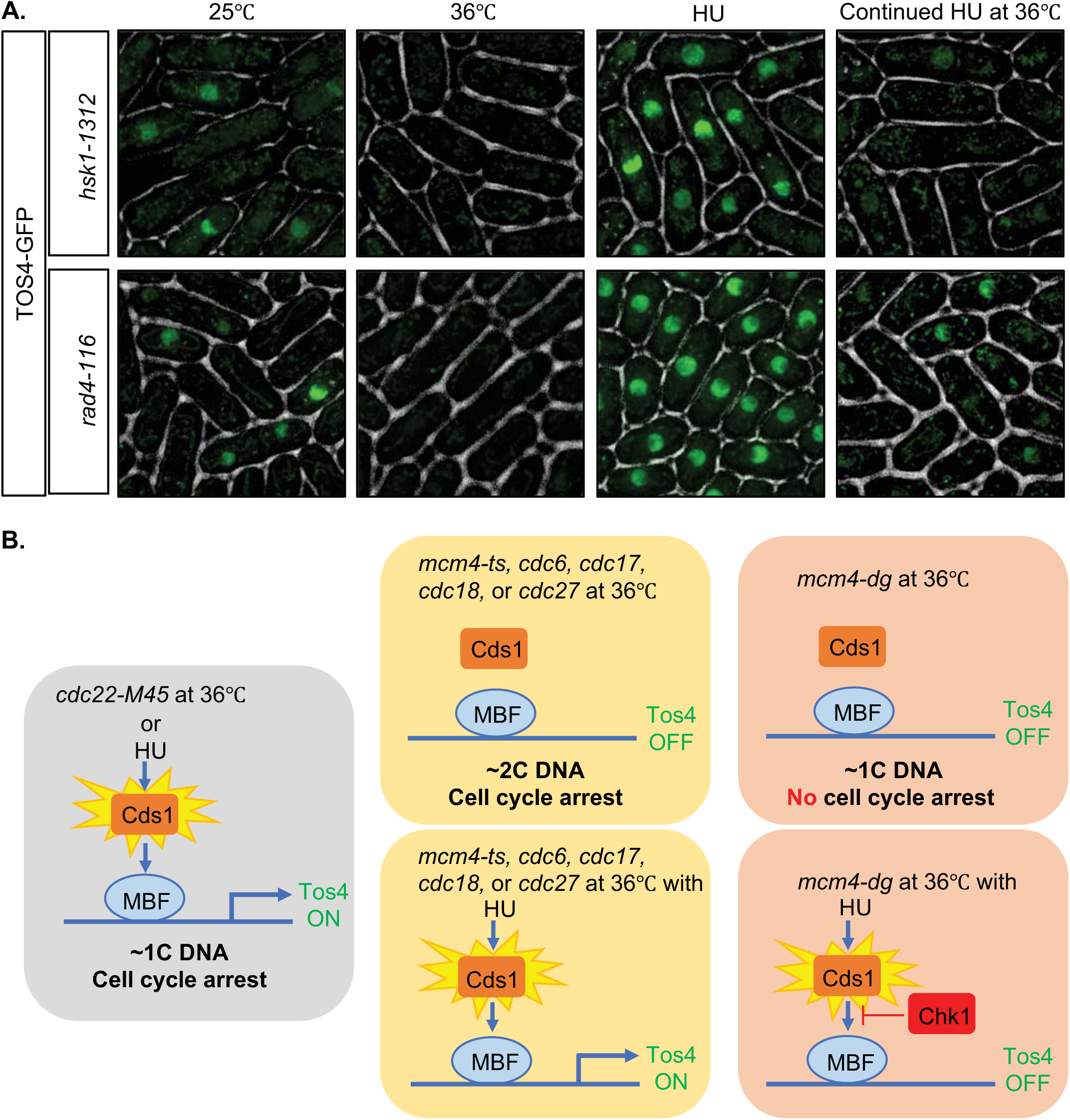
Replication mutants that bypass cell cycle arrest lack nuclear Tos4 despite continued Cds1 activation by HU. (**A**) Temperature-sensitive *hsk1-1312* and *rad4-116* cells were imaged for Tos4-GFP in asynchronous culture at 25°C, after 4h at 36°C, after treatment with 12 mM HU at 25°C, or after pre-treatment with HU at 25°C then transfer to 36°C. (**B**) Left, early replication stress (1C DNA) such as HU and *cdc22-M45* activates Cds1 and Tos4 expression is upregulated. Middle, replication mutants that complete most of DNA synthesis (2C DNA) fail to fully activate Cds1 and Tos4 expression is turned off unless in the continued presence of HU. Right, early replication mutant *mcm4-dg* (1C DNA) that enters mitosis fail to activate Cds1 and turn on Tos4 expression even in the presence of HU, in a Chk1-dependent manner.

## DISCUSSION

Using an imaging-based approach, we demonstrate that nuclear Tos4 accumulation marks early S phase stage independent of DNA content, and allows us to identify three distinctive types of temperature sensitive S phase mutants (Figure 8B): Class 1 mutants arrest replication with a 1C DNA content that results in nuclear Tos4 accumulation; Class 2 are late S/G2 phase arrest mutants (2C DNA content) that lack nuclear Tos4 unless Cds1 is activated by the on-going presence of HU; Class 3 are early replication mutants (1C DNA) that fail to arrest the cell cycle and continue into mitosis, but lack nuclear Tos4 and cannot maintain Tos4 in the nucleus at the restrictive temperature even in the presence of HU.

These phenotypes may be partly distinguished by checkpoint activation. Tos4 is one of many genes whose expression is induced during G1-S transition and repressed in G2 in unperturbed cells, due to oscillation of the MBF transcription factor (Oliveira *et al.* 2012). The activated replication checkpoint kinase Cds1 maintains high levels of G1-S transcription of these genes by preserving MBF activity (reviewed in (Bertoli *et al.* 2013)). Our data show that fission yeast requires active Cds1 to maintain Tos4 accumulation in cells arrested by HU-treatment or by *cdc22-M45* (Fig.2); in budding yeast, the Rad53 checkpoint kinase has a similar effect (Oliveira *et al.* 2012). Lack of nuclear Tos4 accumulation in *cds1*-deleted cells is consistent with transcription and protein levels of Tos4 being similarly decreased in *S. pombe* corresponding to loss of MBF activity. Temperature-sensitive replication mutants that arrest with a nearly 2C DNA content (*mcm4-ts, cdc18-K46, cdc17-M75, cdc17-K42*, *cdc6-23, cdc6-ts2, cdc27-K3*) do not retain nuclear Tos4 at restrictive temperature (Figure 3), consistent with a failure to activate Cds1 (Lindsay *et al.* 1998). Conversely, at least some of these late mutants are known to activate Chk1 in response to double strand breaks, which is required for their arrest (Coxon *et al.* 1992; Forsburg and Nurse 1994; Liang *et al.* 1999; Bailis *et al.* 2008; Yin *et al.* 2008). The loss of Tos4 in these conditions is consistent with a previous study showing that Chk1 inhibits MBF (Ivanova *et al.* 2013). Thus, the two checkpoints have opposite effects on MBF. We conclude that the distinction in Tos4 accumulation in the arrested class 1 and class 2 mutants may represent which checkpoint is active, and the corresponding effect on Tos4 gene expression via MBF (upregulated by Cds1, and downregulated by Chk1).

The third class of mutants *mcm4-dg*, *hsk1-1312* and *rad4-116* have severe defects in DNA synthesis but nonetheless continue into mitosis without a cell cycle arrest, indicating that they have not activated either checkpoint, despite unreplicated DNA and evidence for DNA damage (Saka *et al.* 1997; Snaith *et al.* 2000; Lindner *et al.* 2002; Sabatinos *et al.* 2015). Similarly, they do not maintain nuclear Tos4 at the restrictive temperature, which we conclude may reflect their ongoing cell cycle progression. Consistent with this, we do not observe Tos4 accumulating in *chk1∆ mcm4-dg* at the restrictive temperature.

Interestingly, however, these class 3 mutants also fail to retain Tos4 if shifted to the restrictive temperature in the continued presence of HU, despite the absence of DNA synthesis (Figure 7A, 8A). Our previous data provide some insight into this difference. We showed that *mcm4-dg* cells at the restrictive temperature do not activate Chk1 (Sabatinos *et al.* 2015), and do not show evidence for double strand breaks as measure by accumulation of H2A(X) phosphorylation (Bailis *et al.* 2008). However, if we shift *mcm4-dg* mutants to 36°C in the ongoing presence of HU, they do accumulate H2A(X) phosophorylation (Bailis *et al.* 2008), which suggests they have a different form of disruption or fork collapse than observed in the absence of HU. DSBs activate the Chk1 damage kinase, which we predict should repress the MBF and result in the loss of nuclearTos4. Consistent with this, we observe that a *chk1∆ mcm4-dg* double mutant maintains nuclear Tos4 in HU at the restrictive temperature. (We were unable to determine the effect in *hsk1-1312* and *rad4-116* double mutants because *chk1∆* has synthetic lethality or a growth defect in these backgrounds (Walworth *et al.* 1993; McFarlane *et al.* 1997; Snaith *et al.* 2000; Taricani and Wang 2006). Thus, we conclude that the failure to maintain Tos4 in HU at 36°C for *mcm4-dg* reflects activation of Chk1, which is not the case in the absence of HU. Whether this accounts for the response of *rad4*, which is required for Chk1 activation (Furuya *et al.* 2004) remains to be determined.

Temperature-sensitive APC mutants (*cut9-665*, *cut4-533*, and *nuc2-663)* did not stabilize nuclear Tos4 when released from HU to the restrictive temperature, suggesting Tos4 may not be an APC target in fission yeast. However, Ste9, a WD-repeat protein homologous to budding yeast Cdh1, activates APC and promotes degradation of mitotic cyclins (Kitamura *et al.* 1998; Blanco *et al.* 2000). Cds1 phosphorylates and inhibits Ste9 to protect the MBF activator Rep2 from degradation (Chu *et al.* 2009), and it is possible that Cds1-dependent accumulation of nuclear Tos4 may require Ste9 inhibition.

Thus, the presence of Tos4 in mutant backgrounds at restrictive conditions may be more precisely a measure for different pathways of checkpoint activation rather than position within S phase. Importantly, cells apparently blocked nominally in late S phase may actually be in G2 phase, consistent with recent evidence (Kelly and Callegari 2019). This raises the possibility that the distinction between S phase and G2 actually depends upon the ability to activate the damage checkpoint. Using various fluorescently-tagged cell cycle dependent proteins in combination with other cell cycle and checkpoint mutants will help elucidate whether or not there are distinctive changes that distinguish late S phase and G2 phase.

## ACKNOWLEDGEMENTS

We thank Ji-Ping Yuan for technical support, and Marc Green for initial studies on Tos4. This work supported by NIH R35-GM118109 (SLF).

