## Supplemental Figure 1 for "Checkpoint regulation of nuclear Tos4 defines S phase arrest in fission yeast"

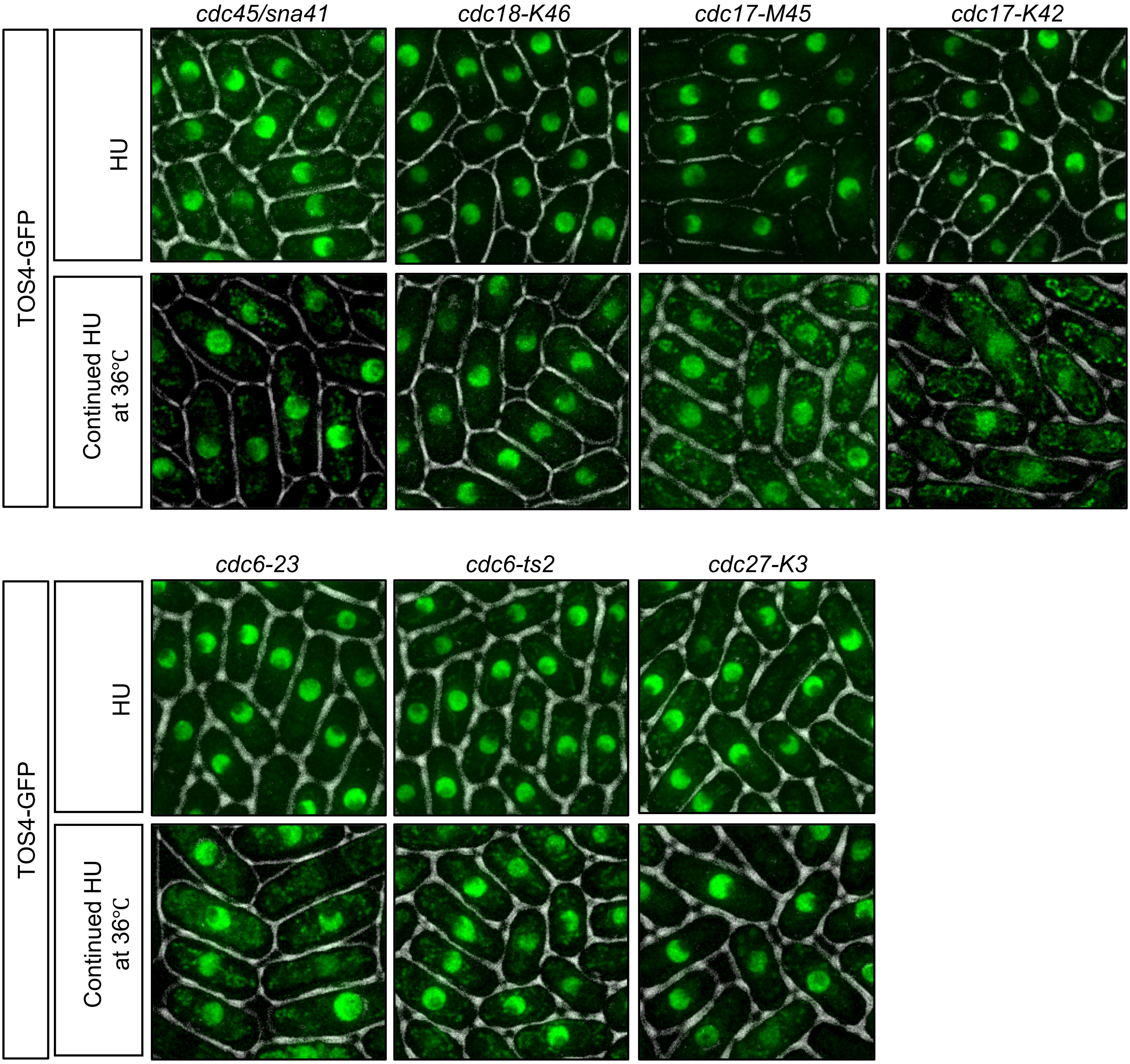


**Figure S1**. Temperature-sensitive replication mutants *cdc45/sna41,* MCM loader mutant (*cdc18-K46*), ligase mutants (*cdc17-M45* and *cdc17-K42*), and polymerase delta mutants *(cdc6-23, cdc6-ts2*, and *cdc27-K3*) were imaged for Tos4-GFP after treatment with 12 mM HU at 25℃, or after pre-treatment with HU at 25℃ then transfer to 36℃.
